# Reproducibility and reusability limitations in Regulatory Circuits: analysis and solutions

**DOI:** 10.1101/2021.08.02.454723

**Authors:** Marine Louarn, Anne Siegel, Thierry Fest, Olivier Dameron, Fabrice Chatonnet

## Abstract

The Regulatory Circuits project is among the most recent and the most complete attempts to identify cell-type specific regulatory networks in Human. It is one of the largest efforts of public genomics data integration, based on data from the major consortia FANTOM5, ENCODE and Roadmap Epigenomics. This project is a main provider of biological data, cited more than 224 times (Google Scholar) and its resulting networks were used in at least 42 other articles.

For such a general resource, reproducibility of both the outputs (regulation networks) and methods (data integration pipeline) is a major issue, since biological data are updated regularly. In addition, users may want to introduce new data into the Regulatory Circuits framework to provide networks about previously uncharacterized cell types or to add information about specific regulators, which require to re-execute the whole pipeline on the new data.

In this article, we analyze the various factors limiting reproducibility of the Regulatory Circuits data and methods. Starting from a factual description of our understanding of the methods used in Regulatory Circuits, our contribution is two-fold: we propose (1) a characterization of the different levels of reusability, reproducibility and conceptual issues in the original workflow and (2) a new implementation of the workflow ensuring its consistency with the published description and allowing for an easier reuse and reproduction of the published outputs. Both are applicable beyond the case of Regulatory Circuits.

## 1. Introduction

Life science, and in particular health, is a major data producer: for example the US healthcare system as reached 150 exabytes in 2010 (1), and this trend is expected to increase over the next decade (2). In addition to the data quantity challenge data heterogeneity is a second challenge. In (3) the authors performed a review of different fields of data production in health: from genomics, proteomics, metabolomics, to imaging, clinical diagnosis, patient history and the recent addition of personal devices information. Moreover, we are confronted to the rise of multi-omics solutions (Genomics, Epigenomics, Proteomics, Transcriptomics, Metabolomics, etc…) in health and disease-related research (4, 5). Each dataset’s size often requires to split the data into several files, which adds another layer of integration and makes global analysis all the more complicated (6).

Even in the specific domain of gene regulatory networks, the data are large and heterogeneous. To understand the regulation in a given biological context, one needs to perform diverse types of experiment spanning spanning whole genomes, currently made possible by the recent advent of high throughput sequencing. A gene regulation network encompasses the interactions either between regulators, or between a regulator and other entities in a cell to control genes expression. For gene regulatory networks, regulators are specialized proteins called transcription factors (TF) which interact with DNA, the molecular support of genetic information. At the DNA level, a TF will bind to a definite sequence (called a binding site) in a specific regulatory region, which should be in an opened 3D conformation to allow the regulation (7, 8), and which can be located close or far from its target gene (9). This binding event will then initiate a cascade of molecular events eventually leading to regulation (induction or inhibition) of the target gene’s expression.

A better characterization of gene regulatory networks allows a better understanding of major processes such as cell differentiation (how to obtain one or several effective cell types from a common progenitor cell), cell identity (how gene expression is used to define a specific cell type) and cell transformation (how altered gene expression can lead to cell death or cancer) (10). Many gene regulatory network inference methods have been published, but few of them apply to human data, and even fewer are able to take into account inputs from gene expressions, regulatory regions activities and transcription factor binding sites (11, 12). Among the latter, the *Regulatory Circuits* project^1^ (13) is one of the largest effort of genomics data integration in human cells. It consists of several analyses on heterogeneous and multi-layer “omics” data on 394 human cell lines and primary cells from tissues. This project is a major provider of biological data, cited more than 224 times (Google Scholar – 2021-06-30). Its resulting networks were used in at least 42 other articles.

Despite being such a fundamental resource, this project has not been updated since its publication in 2016. The *Regulatory Circuits* website gives access to unstructured, disconnected and diversely formatted tabulated files related either to source biological data (FANTOM5 data, genes and regions genomic coordinates, TFs binding sites occurrences… divided in 26 files), to computation intermediate results (59 files), or to the results of *in silico* integrative analyses (394 files, one for each network). The outputs of the project are only available as text files and no functional pipeline or software is provided to perform new computations. This has huge impact on i) the reproducibility of the results, ii) their maintenance as they will need to be updated when newer or additional data sources are released and iii) their reuse for advancing other studies (which was the reason these results were generated in the first place).

## 2. What is *Regulatory Circuits*: biological model, input / output data and computational concept

Figure 1 presents the biological principles behind regulatory circuits. The interaction between a TF and a gene is determined as the ability of the TF to bind in a region close the gene, and therefore to regulate (i.e. modify) its expression. Two types of regulatory regions are described, depending on their distance to the gene locus (its location on DNA): promoters are immediately close to the gene and usually encompass its transcription start site (TSS) whereas enhancers are acting at long range, from several to hundreds of kbs. Genes can be described by functional units called transcripts (or isoforms), which may be different for the same gene, depending on alternative TSS usage or which exons (the informative or “coding” pieces of a gene) are used to build a specific transcript. In *Regulatory Circuits*, the relations are computed between TF and transcripts, either considering the promoters or the enhancers and then the relations are grouped by TF and genes, giving unique TF-gene relations (see below).

**Figure 1:**
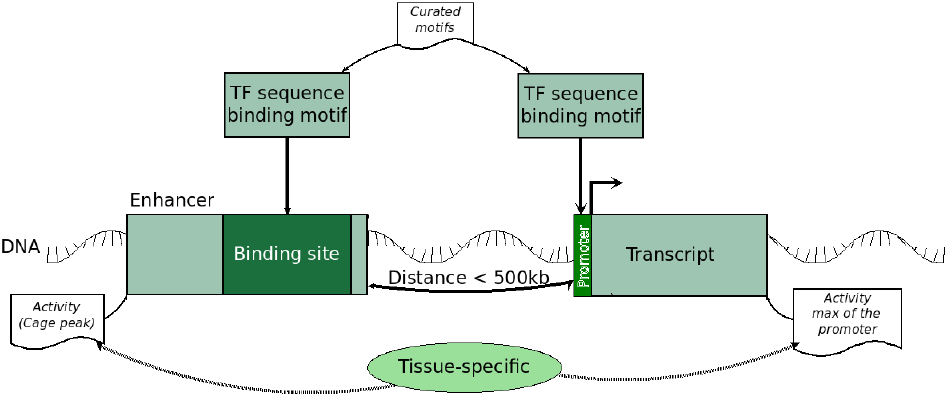
Biological principles behind *Regulatory Circuits*. A transcription factor protein binds to a DNA region called enhancer or promoter depending on its distance to a gene, which regulates the gene expression. Characterizing the relation between a transcription factor and a gene requires to measure the quantity of transcription factor, the opening of the 3D structure of DNA in the binding site, and the observed quantity of the gene’s transcript.

The entry data used in *Regulatory Circuits* comes from several independent projects: FANTOM5 (14) for measuring transcription or regulatory activity (using CAGE-peak data on the regulatory regions, 808 different samples having information for both enhancers and promoters), ENCODE (15) for the prediction of transcription factor binding sites, and GTex (16) and Roadmap Epigenomics (17) for validation data.

*Regulatory Circuits* computes networks that are not derived from a statistical analysis of biological measurements but based on a set of computed correlations between regulatory regions activities, gene expressions, and curated and scored TF binding sites.

Published datasets encompass both input (raw data) and intermediary (authors-processed) data files, in the form of tabulation or comma-delimited data files with various formats and contents.

The output of the Regulatory Circuits study is a set of 394 scored tissue-specific regulatory interaction networks that can be explored through text files, representing 9.1 GB. Each file basically contains three columns: one for the TF, one for the genes and one for the computed score of the relation.

## 3. What is *Regulatory Circuits*: Detailed workflow

Figure 2 presents an overview of the *Regulatory Circuits* workflow, each step is described in the following subsections, as we understood it from a careful revision of the provided methods and supplementary data.

**Figure 2:**
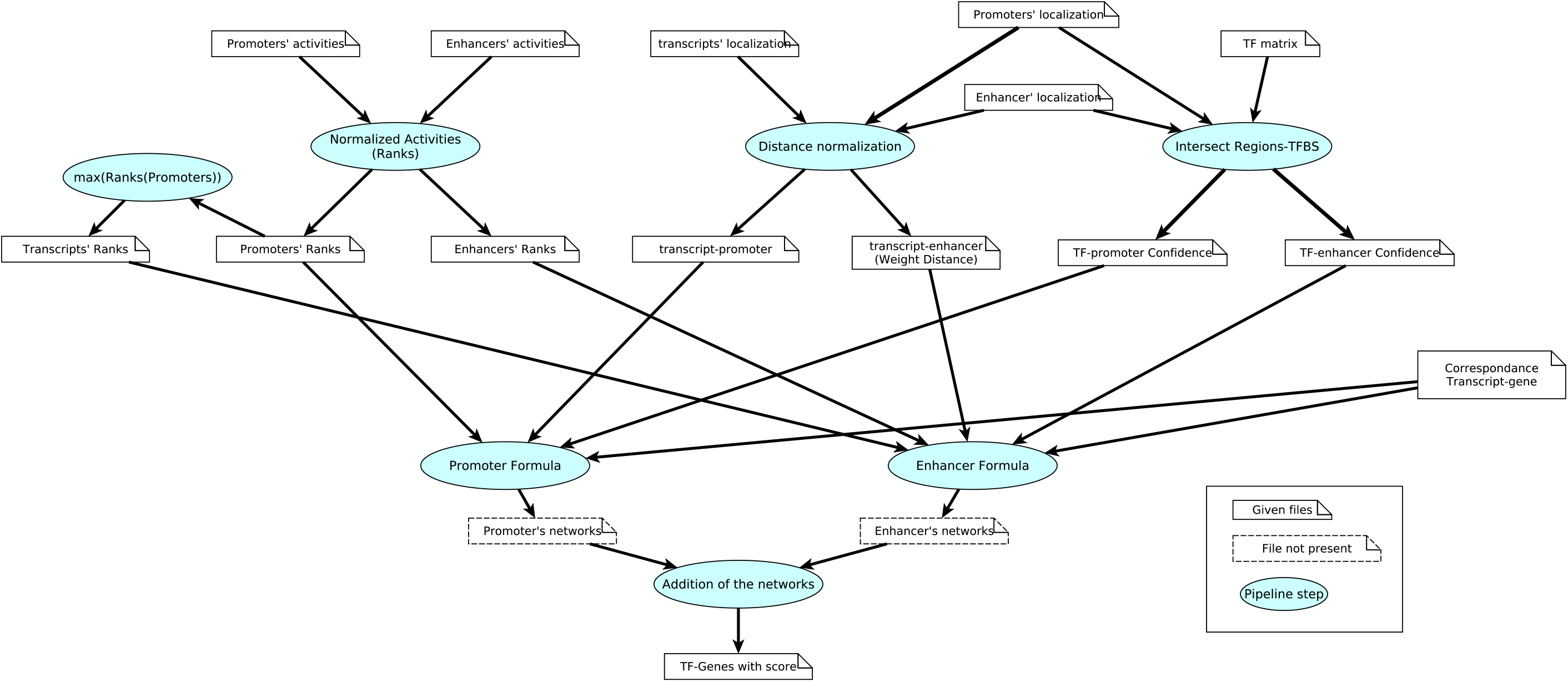
*Regulatory Circuits* global workflow and its different steps.

### 3.1. Global formula

The score (*w*_*ij*_(*S*)) of a relation between the TF (*i*) and the transcript (*j*) in the sample *S* is based on the distance weight (*d*_*jk*_) between the transcript and the regulatory region (*k*), the confidence score of the binding site of the TF in the given region (*c*_*ik*_), the normalized activity of the region (*x*_*k*_) and the normalized activity of the transcript (*y*_*j*_). Giving the following formula:

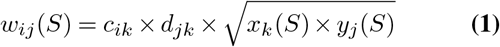

Note that according to Regulatory Circuits, 0 ≤ *d*_*jk*_ ≤ 1 and that *d*_*jk*_ decreases as the distance between the regulatory region and the gene increases, so *d*_*jk*_ is a normalized weight that behaves like the inverse of the distance (see Section 3.4). For Promoters, the distance weight is normalized to 1 -as the promoter is adjacent to the transcript- and the transcript activity is taken as its promoter activity. The previous formula is then reduced to :

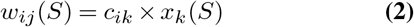

### 3.2. Normalized expression or activity of regions and transcripts

As mentioned in the previous section, the activities of the different elements are normalized.

The choice of the normalization is important in such workflows as the value directly influences the score of the relation between the TF and Genes in the resulting networks, a shown by the equations 1 and 2.

In *Regulatory Circuits* the authors normalize the activities of the different regions as a score between 0 and 1 computed element by element but do not provide the normalization function.

> The weight of promoter-gene edges was defined as the normalized activity level of the promoter across all samples (normalization was done per regulatory element because expression levels of diverse enhancers and promoters might not be on the same scale). Thus, if the promoter is not active in a given cell type, the edge weight is 0 (i.e., the edge is not present), and if the promoter is maximally active, the edge weight equals 1. (13)

For the transcripts, the activity was based upon their promoters:

> The activity level of isoforms was defined as the maximum activity level of their promoters (which are usually few—the majority of isoforms have only one or two alternative promoters). (13)

In the article, the normalization function used is not further described.

### 3.3. Confidence score of the TF binding sites

In *Regulatory Circuits*, the authors consider 662 TF and their binding motifs (position weight matrices) in the genome. They used a curated collection of matrices and assigned a confidence score to each binding site based on the conservation across mammals using the works by Kheradpour et al. (18–20).

The binding sites were looked for in a 400 bp upstream to 50 bp downstream window of the considered promoter, and limited to the actual chromosomal coordinates of the enhancers. If several binding sites of a same TF were found in the same regulatory region, the maximum of their confidence scores was assigned to the TF-region relation, regardless of the binding site location inside the region. The scores and relations between TF and regions are compiled in the *tf- - -promoter. prec90*.*txt* and *tf- - -enhancer. prec90*.*txt* files provided in *Regulatory Circuits* supplementary data.

### 3.4. Distance weight of the regions

The weighting function for the distance only applies to enhancers For promoters the weight of the distance is set to 1.

To normalize the distance between enhancer and transcript, the authors used cis-eQTLs from RegulomeDB (21) and computed their distance to the TSSs of target genes. The weigh function is defined using a local polynomial regression fitting for the range 1 kb to 500 kb (in either direction from the transcript), where 1 kb was normalized to 1 and 500 kb to 0. This is then applied to the enhancers considered in the workflow.

This means that all enhancers further than 500 kb of the transcript were not considered for the remainder of the computation. The formula to compute the distance weight was provided in the workflow description.

### 3.5. From individual relations to networks

Up until this point all relations are computed using the transcripts instead of genes. To determine the TF-genes relations, all the relations between a TF and transcripts of a same gene are merged into one and its score is the maximum of the scores of the TF-transcripts relations.

Similarly, if a TF-gene relation exists using several promoters or enhancers, the relation is kept using the maximal score computed, for each type of regulatory region.

> For each pair of edges forming a chain that connects a TF to a promoter to an isoform […]. If several redundant edges between the same TF and gene were found (via different promoters or isoforms), they were merged and the maximum edge weight was retained. A separate TF-gene network encapsulating all regulatory interactions via enhancers was created using the same approach. (13)

At this point, the *Regulatory Circuits* workflow gives two ways of calculating a TF-gene relation and its associated score: one using enhancers and the second using promoters. In the resulting networks this distinction is not made and both ways of computing have been combined to obtain the final score of the TF-gene relation:

> Both TF-gene networks thus had edge weights ranging from 0 (absent edge) to 1 (highest confidence), which were added to form a combined TF-gene network including evidence from both promoters and enhancers. (13)

## 4. Issues with *Regulatory Circuits*

While trying to run and understand the *Regulatory Circuits* workflow we encountered multiple setbacks as summarized in Figure 3.

**Figure 3:**
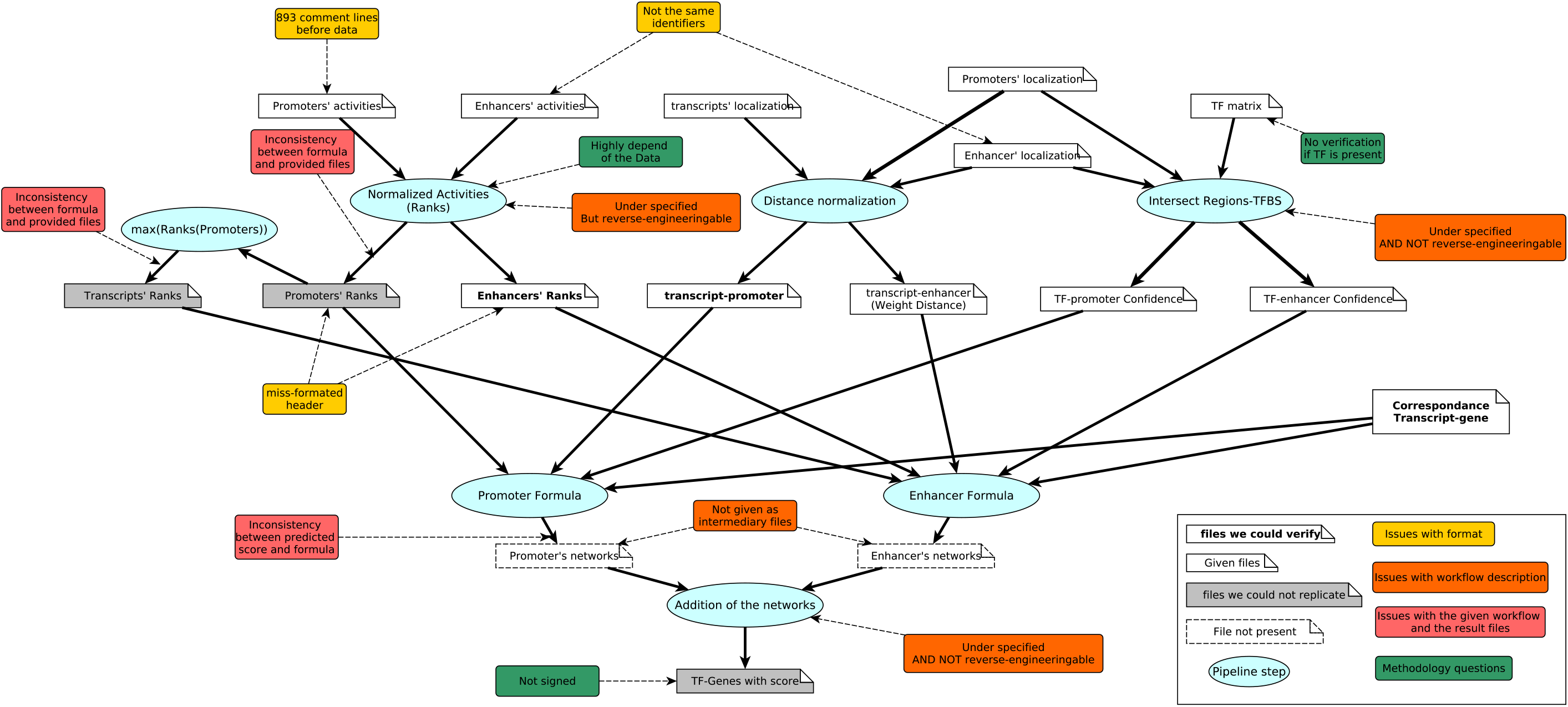
*Regulatory Circuits* global workflow as presented in Figure 2 with the addition of the point of contention: either issues in the workflow or remaining doubt on the step.

Among them, we identified difficulties with understanding the overall dependencies between the files as well as their structure and content (Section 4.1). Moreover, the scripts provided by *Regulatory Circuits* could not be used on the project’s original files nor on other files (Section 4.2). We also noted that some of the intermediary files were not provided (Section 4.3), which was a problem because we could not validate the steps producing them (Section 4.4), i.e. compare what we get when we re-implement the steps with the original results. More generally, we identified problems with *Regulatory Circuits* method that will hamper the method’s reusability in similar biological contexts (Section 4.5). Eventually, the general implementation of *Regulatory Circuits* relies on *ad hoc* technical choices that also limit both the method and final results reusability (Section 4.6) and are not on par with FAIR recommendations. As shown in Figure 3, all these limitations occur multiple times over the *Regulatory Circuits* workflow.

### 4.1. Understanding the files

We analyzed the 21 input or intermediary files provided by *Regulatory Circuits*. Table 1 summarizes the files characteristics.

**Table 1:** Regulatory Circuits files’ review

| type of file | file name | format | header lines | missformatted header | comment lines | data lines | nb columns | Column(s) with ID | ID format | Entities | Source | content |
| --- | --- | --- | --- | --- | --- | --- | --- | --- | --- | --- | --- | --- |
| Input Data for network inference | *hg19_permissive_enhancers_expression_rle_tpm.csv* | csv (,) | 1 |  | 0 | 43011 | 809 | 1 | chr:start-end | enhancer | [1] | [a] |
|  | *permissive_enhancers.bed* | bed12 (tab-delim) | 1 |  | 0 | 43011 | 12 | 4 | chr:start-end | enhancer | [1] | [b] |
|  | *robust_enhancers.bed* | bed12 (tab-delim) | 1 |  | 0 | 38554 | 12 | 4 | chr:start-end | enhancer | [1] | [b] |
|  | *hg19.cage_peak_tpm.osc.txt* | tab-delim | 3 |  | 893 | 184827 | 890 | 1 | chr:start-end,strand | promoter | [2] | [a] |
|  | *hg19.cage_peak_coord_robust.bed* | bed12 (tab-delim) | 0 |  | 0 | 184827 | 12 | 4 | chr:start-end,strand | promoter | [2] | [b] |
|  | *gene_coord.bed* | bed6 (tab-delim) | 0 |  | 0 | 19125 | 6 | 4 | GENE_SYMBOL | gene | [3] | [b] |
|  | *gene_ids.txt* | tab-delim | 1 |  | 0 | 19125 | 3 | 1 | ENSG000000000000 | gene | [3] | [c] |
|  |  |  |  |  |  |  |  | 2 | GENE_SYMBOL | gene |  |  |
|  |  |  |  |  |  |  |  | 3 | EntrezID | gene |  |  |
|  | *mhc_genes.txt* | tab-delim | 1 |  | 0 | 184 | 1 | 1 | GENE_SYMBOL | gene | [3] | [c] |
|  | *transcript_coord.bed* | bed6 (tab-delim) | 0 |  | 0 | 53449 | 6 | 4 | GENE_SYMBOL-000 | transcript | [3] | [b] |
|  | *transcript—gene.txt* | tab-delim | 1 |  | 0 | 53449 | 4 | 1 | GENE_SYMBOL-000 | transcript | [3] | [c] |
|  |  |  |  |  |  |  |  | 2 | GENE_SYMBOL | gene |  |  |
|  |  |  |  |  |  |  |  | 3 | ENSG000000000000 | transcript |  |  |
|  |  |  |  |  |  |  |  | 4 | ENST000000000000 | gene |  |  |
|  | *tss_coord.bed* | bed6 (tab-delim) | 0 |  | 0 | 53449 | 6 | 4 | GENE_SYMBOL-000 | gene | [3] | [b] |
|  | *motif_defs.txt* | space-delim | 0 |  | 1772 | N/A | N/A | N/A | N/A |  | [4] | [g] |
|  | *motif_instances.bed* | bed6 (tab-delim) | 0 |  | 0 | 124358159 | 6 | 4 | TF_0 | TF | [4] | [b] |
|  | *tf_motif_ids.txt* | tab-delim | 1 |  | 0 | 1792 | 3 | 1 | TF | TF | [4] | [g] |
|  |  |  |  |  |  |  |  | 2 | TF_0 | TF |  |  |
| Intermediary files | *enhancer_expr.rank.txt* | tab-delim | 1 | x | 0 | 43011 | 809 | 1 | e@chr:start..end | enhancer | [5]* | [d] |
|  | *enhancer—transcript.prec90.txt* | tab-delim | 1 |  | 0 | 950513 | 5 | 1 | e@chr:start..end | enhancer | [5]* | [e] |
|  |  |  |  |  |  |  |  | 2 | GENE_SYMBOL-000 | transcript |  |  |
|  |  |  |  |  |  |  |  | 5 | GENE_SYMBOL | gene |  |  |
|  | *promoter_expr.rank.prec90.txt* | tab-delim | 1 | x | 0 | 59126 | 809 | 1 | p@chr:start..end,strand | promoter | [5]* | [d] |
|  | *promoter—transcript.prec90.txt* | tab-delim | 1 |  | 0 | 123440 | 4 | 1 | p@chr:start..end,strand | promoter | [5]* | [e] |
|  |  |  |  |  |  |  |  | 2 | GENE_SYMBOL-000 | transcript |  |  |
|  |  |  |  |  |  |  |  | 4 | GENE_SYMBOL | gene |  |  |
|  | *tf—enhancer.prec90.txt* | tab-delim | 1 |  | 0 | 524816 | 3 | 1 | TF | TF | [5]* | [f] |
|  |  |  |  |  |  |  |  | 2 | e@chr:start..end | enhancer |  |  |
|  | *tf—promoter.prec90.txt* | tab-delim | 1 |  | 0 | 1169797 | 3 | 1 | TF | TF | [5]* | [f] |
|  |  |  |  |  |  |  |  | 2 | p@chr:start..end,strand | promoter |  |  |
|  | *transcript_expr.rank.prec90.txt* | tab-delim | 1 | x | 0 | 43352 | 809 | 1 | GENE_SYMBOL-000 | transcript | [5]* | [d] |

Sources: [1]http://enhancer.binf.ku.dk/Pre-defined_tracks.html, [2]http://fantom.gsc.riken.jp/5/datafiles/latest/extra/CAGE_peaks/, [3] Ensembl bioma, [4] Pouya Kheradpour & [5]\*
Regulatory Circuits: auto produced
Type of content: [a] Normalized activities, [b] Genomic coordinates, [c] Identifier, [d] Rank of normalized activities, [e] Distances, [f] Confidence score & [g] TF motifs

A first setback arose during the inspection of these files. It was related to the difficulty to explore them and to extract the information they contain. Some of the files have non explicit names regarding what they contain (e.g.: *hg19. cage_peak_tpm. osc. txt* which contains the expression of the promoters). All the files are not in the same format: some were comma-separated, others, tabulation-separated. Fifteen of the files had headers, while 6 did not. Moreover, two of the files presenting header had misaligned header to data (missing column). Two of the 21 files start with some comments before the data, ranging up to 1700 lines.

It was also difficult to link the data between two files as the identifiers referring to the same entity were not consistent among the files (e.g.: chr:start-end in file *permissive _enhancers. bed* corresponds to *e@chr:start-end* in *tf- - - enhancer. prec90. txt*) as shown in Table 1 column “ID format”.

### 4.2. *Regulatory Circuits* scripts are not usable

The computational scripts and algorithms provided as resources are limited to the considered data-set. The given implementation was made using Java but is unusable as such as it lacks explanation on the input files necessary and on how to run it. Furthermore, there is a lack of documentation on the implementation, the wiki is still under construction^2^ and have not been updated since 2015. The project website^3^ has not been updated and the last news on the project were from august 2016. As of June 2021 the website domain expired and the site is therefore unavailable.

Moreover, we contacted the Regulatory Circuits’ authors in July 2018 with solicitations about the methodology but we did not get any feedback for the moment.

### 4.3. Intermediary files not present

While most of the pre-processed data are present in the download folder, the authors do not give any access to three intermediary steps of the workflow. The intermediary networks obtained by different type of regulatory regions (promoters or enhancers) are not provided, nor are the intermediary sample-specific networks. This led to issues while trying to reverse-engineer the *Regulatory Circuits* workflow as we could not check the computed intermediary scores before the last step.

### 4.4. *Regulatory Circuits* methodology could not be reproduced

#### Rank

We ran into several issues with steps of the workflow while trying to understand it: the normalization function used for the expression, the weight function for fitting the enhancer-gene distance as well as the notion of ’addition’ of enhancer and promoter network (“Both TF-gene networks[…] were added to form a combined TF-gene network”) were all under-specified, and some steps were not performed as described: the rank of the promoter was not computed element by element. For some parts of the pipeline, we were nevertheless able to reverse-engineer part or all of the published methodology.

We also found inconsistencies between the described methodology and the result files. For the ranks of the promoters the formula was not applied element by element as described but on all the promoters of a same transcript at the same time.

Using reverse engineering, we managed to find out for the enhancer that the normalization is an application of the rank function, this information was given in the name of the transformed expression files: *enhancer_exp.rank.txt, promoter_expr.rank.prec90.txt* and *transcript_expr. rank.prec90.txt*.

For each enhancer the activities in the different samples are ordered from the least expressed to the most expressed. Several samples can have no expression and their ranks are set to 0. The other samples ranks are computed as: Position in which they appear after being ordered divided by the number of samples expressed (expression strictly > 0). For an enhancer, the most expressed samples are therefore normalized to 1.

For promoters, we realized when reverse-engineering the ranks that they were not computed element by element (i.e. promoter by promoter) in this step. Instead all promoters were grouped based on the transcript they were preceding, and the rank function was applied to the expression data of all these promoters. Therefore, according to the *Regulatory Circuits* method, if a transcript is preceded by n promoters, the samples ranks should have been computed separately for each promoter on the 808 samples (therefore, for each promoter, the sample ranks span [0; 1]). Instead, we observed that the samples ranks were computed globally on the union of the n promoters (therefore, for one promoter, the sample ranks may span a sub-interval of [0; 1]). Figure 4 illustrates the differences between the expected and observed promoter ranks for a transcript with eight promoters.

**Figure 4:**
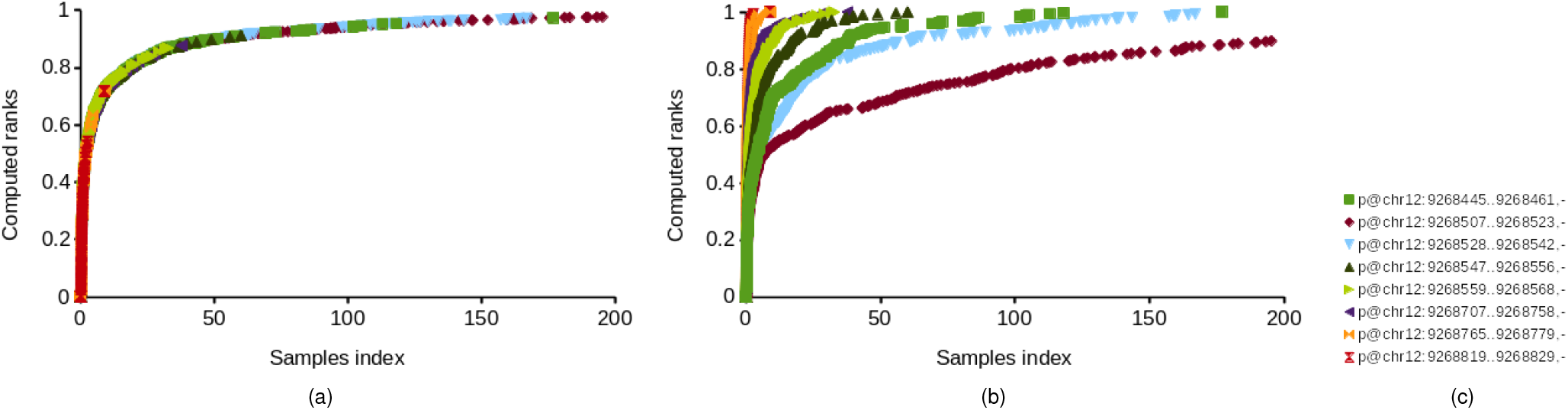
Difference between the promoter ranks for a transcript with eight promoters observed in intermediary files (4a) and the promoter ranks computed according to the method detailed in (13) (4b). 4c list the eight promoters of the transcript. Please note that in 4a, all ranks are aligned on one curve denoting the use of a union of promoters to compute the transcript ranks, while on 4b each promoter has its own curve, ranging from 0 to 1.

For the each transcript, its rank for a given sample was supposed to be the maximum rank of its different promoters in this sample, but it was not what we found in the intermediary files. The transcripts’ ranks were computed as the means of its promoters ranks weighted by a different constant for each transcript. Therefore we could not undoubtedly compute the transcript scores and abandoned our retro-engineering process. This is one of the reasons why we chose to re-compute the entire networks (see Section 5).

#### Distance relation

Using the pre-computed file *enhancer- - -transcript. prec90.txt* provided by *Regulatory Circuits* that indicate the weighted distance, we were able to compute an approximation of the polynomial regression and to apply it.

#### Overall scores

When looking into the resulting networks we found relations scored to values superior to 1. The addition operation described between the promoter weight and the enhancer weight, was therefore not a max as for the enhancer - or promoter - weights across the samples constituting the tissue. We were not able to compare the intermediary weight to the original one founds (as no files were given) and therefore could not confirm the *addition* used.

When computed using a max relation between promoter and enhancer weight, the overall scores we found were superior to the ones found in the original study. Using a sum relation between those weights to compute the score would have made the difference even greater as our score would have increased.

#### Overall networks

Eventually, we found discrepancies between the potential relations given in the output files and what could be computed in simple case (mostly relations provided by *Regulatory Circuits* that we could not reproduce, and occasionally relations that we predicted but did not appear in *Regulatory Circuits* final results). For example, in the Myeloma cell line, the TF INSM1 could only regulate the gene *AMER1* through one specific promoter and with no enhancer involved. The relation is scored (4.21371298E-03) in the corresponding output file, but the confidence score of the TF in this promoter is given at 0 and applying formula 2 would also yield a 0 global score. Figure 5 shows the disparity between the relations found in the result files given by *Regulatory Circuits* (union of all TF-gene relations across all networks) and the relations we found by computing the relations between entities across all files (following the methodology and simple bash commands, like *join*, see section 5).

**Figure 5:**
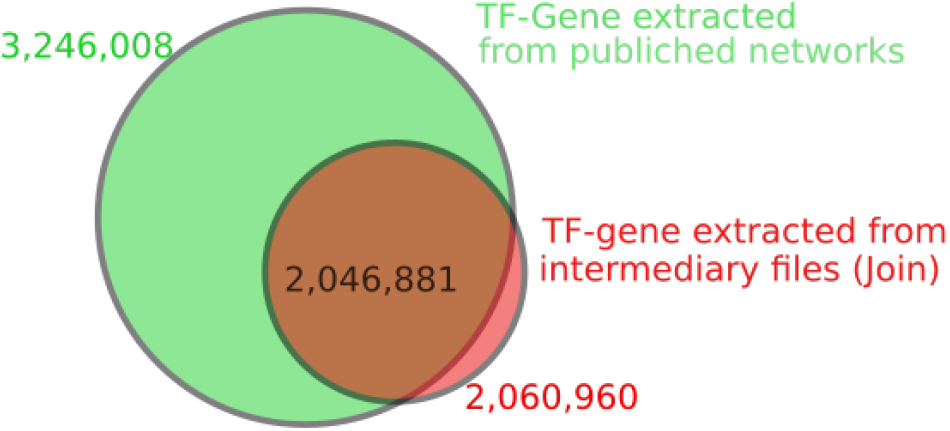
Number of unique TF-genes relations found in results network (in green) and number of relations found while using the files and the described method (red). The relations in green but not in red can be explained by the the attribution of a positive score to relations which should have been scored to 0, and by some minor changes in network topology. The relations in red but not in green could be explained by relations wrongly scored to 0, probably because of the discrepancies between the described and the actually used method, e.g. concerning the ranks.

### 4.5. Conceptual issues

*Regulatory Circuits* stops at the tissue level in the original computed networks, while some users may want to look at finer levels such as samples of a same tissue. The provided method therefore lacks flexibility to have fine-grained / personalized networks.

The way of computing scores seems highly biased toward activation, as the scores are computed by multiplying the maximums of several parameters (activities, confidence scores, distance scores…). For the tissue-specific networks, the relations are therefore unsigned and potential inhibition are either hidden or lost. For the same reasons, genes which are less expressed in one sample or tissue are more susceptible to be excluded from a network, disregarding their potential function (for example, TF coding genes are known to be expressed at low to moderate levels).

Additionally, the rank function used for normalizing the expressions only takes their order into account not the distance: for example, to samples with successive expressions of 0.1 and 0.6, or of 0.6 and 0.7 will have the same distance in Rank (1/nb samples). It is not a fine grain normalization function. Removing or adding samples can drastically change the rank distribution: focusing on a portion of *Regulatory Circuits* means that computed ranks will be different from the ones provided by *Regulatory Circuits*, i.e. the latter are not reproducible. We hypothesized that the ranks of two biologically close samples would be really similar among a lot of other samples, and would be wrongly separated among very few others, questioning the reuse of the *Regulatory Circuits* methodology on limited numbers of samples (a few tenths at most), a common setting in life and medical sciences and in clinical settings.

Another issue with the methodology is that the networks assume the TF to be expressed in the cell type, but the workflow never checks if it is really the case. All relations found are assumed to be applicable but if a TF is not expressed in a specific cell type, then the regulation involving this TF may not exist.

### 4.6. FAIR-related problems with method and results re-usability

The networks’ output format -Text files-makes it impossible to explore and enrich the data by combining them to additional knowledge on entities stored in LOD public databases.

The workflow design makes it difficult both to extend *Regulatory Circuits* by adding new data or updating them, and to reuse only part of *Regulatory Circuits*, as it requires to recompute all the ranks for all the entities and neither the provided programs nor the incompletely specified method allow it. It is also difficult to enrich with new types of information: adding gene expressions would mean defining a new formula for scores, as the promoter can not be approximated by the gene (several promoters of one transcript and several transcripts for one gene).

## 5. Computing *Regulatory Circuits* networks

*Regulatory Circuits* is a general resource on regulatory networks. We wanted to use it on specific cells-types not included in the published networks. To be able to run the *Regulatory Circuits* workflow on new data, we devised a new bash-based implementation to compute the original regulatory networks, using most of the provided input and intermediary files and strictly following the methods as described and understood in Section 3.

### 5.1. A new way of computing *Regulatory Circuits* networks

For running *Regulatory Circuits* the first step was to identify the necessary files and homogenize the identifiers for all entities across the files.

We re-computed the ranks as described in *Regulatory Circuits*. We used the R rank function separately on each regulatory element as the principle is the same for both enhancers and promoters. For the transcripts, we took the rank of their promoter and for each sample kept the highest rank across all promoters. The remainder of the steps of the workflow are computed using the intermediary files of the original study (confidence score, distance, relations between entities, etc.). For the final steps - computing the scores - we applied the formula provided in the paper, and kept the maximum score for a TF-genre relation: the maximum between the score by promoter and by enhancer and the maximum score in all the samples constituting the tissue. All the scripts and necessary files are available at https://gitlab.com/mlouarn/RegulatoryCircuits. Table 2 compares the different steps of the original workflow and the new pipeline of calculating the resulting tissues-specific networks.

**Table 2:**
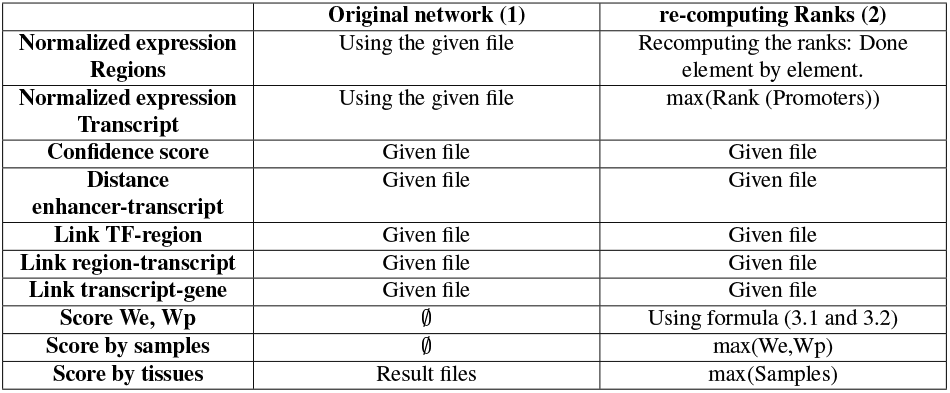
Comparison between the original *Regulatory Circuits* workflow and the new pipeline. We: scores for sample-specific network using only enhancers, Wp: scores for sample-specific network using only promoters

### 5.2. Comparing the original workflow to the new way to compute *Regulatory Circuits* networks

We compared two ways of obtaining *Regulatory Circuits* networks: the original study using directly *Regulatory Circuits* output network files (1) and our bash-based solution only using input files and computing all the steps we could recompute (2). For all tissues and samples, all relations between TFs and genes found in the re-computed method were relations conserved from the original study networks. However, we also identified 1,185,048 TF-genes relations (36.5% of the originally published relations) that were not predicted by our implementation. For all of these relations, at least one of the weights was zero, which resulted in a final score of zero. We have no explanation on why these relations had a strictly positive score in *Regulatory Circuits*.

For an in-depth analysis, the comparison was performed on 12 cells-types: B lymphoblastoid cell line, brain fetal, CD4+ T cells, CD8+ T cells, CD34+ stem cells-adult bone marrow derived, colon adult, colon fetal, epitheloid cancer cell line, pancreas adult, peripheral blood mononuclear cells, small intestine adult and small intestine fetal. We used 12 tissues on which we had RNA-seq data (from Roadmap epigenomic) to run similar validation of the networks as done in the original paper.

#### 5.2.1. Networks topology

In Table 3 we show the variability of the regulatory networks depending on the two strategies. In the computed networks, we lost an average of 50 TFs in the result graphs but retained a similar number of genes and lost nearly half of the relations in the final networks.

**Table 3:** Comparison between the obtained networks on 12 cells types. Our method produce networks that are included in the original *Regulatory Circuits* networks.

| Type of Circuits | Given as Result |  |  | Re-Computed |  |  |
| --- | --- | --- | --- | --- | --- | --- |
|  | min | max | mean | min | max | mean |
| Nb of TF | 643 | 643 | 643 | 593 | 596 | 594 |
| Nb of Genes | 11,911 | 14,850 | 13,067 | 11,881 | 14,812 | 12,933 |
| Nb of Relations | 407,056 | 1,796,098 | 1,042,839 | 239,049 | 1,010,008 | 581,711 |
| % of complete graph | 5.2 | 21.9 | 12.5 | 3.3 | 13.3 | 7.6 |

*Regulatory Circuits* compared the genes regulated in the tissue-specific networks to the expressed genes of the RNA-seq of the Roadmap Epigenomics project. The authors conclude that, as expected, the highly expressed genes were largely (more than 90%) recovered in the produced networks and that the least expressed genes had no regulatory input (less than 10% recovered).

We did the same analysis on the networks computed with the two above-mentioned methods (Figure 7). We found that the re-computed network had slightly lower level of recovery of the genes, but not significantly (Wilcoxon rank sum test between the two series of recall measures for 12 tissues).

**Figure 6:**
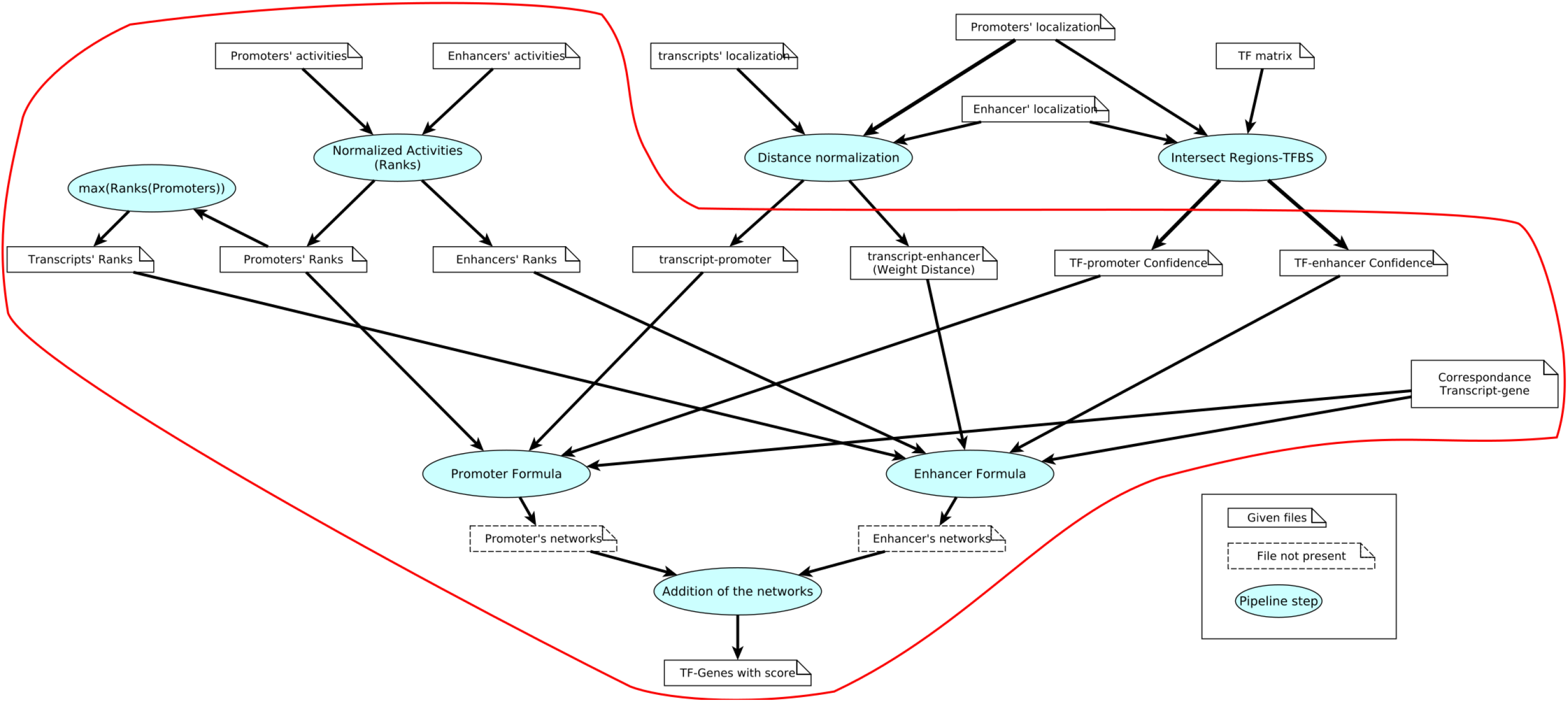
*Regulatory Circuits* global workflow, circled are the files reused and the steps re-computed of the workflow. Circled in red is the re-implementation in which we recompute the ranks.

**Figure 7:**
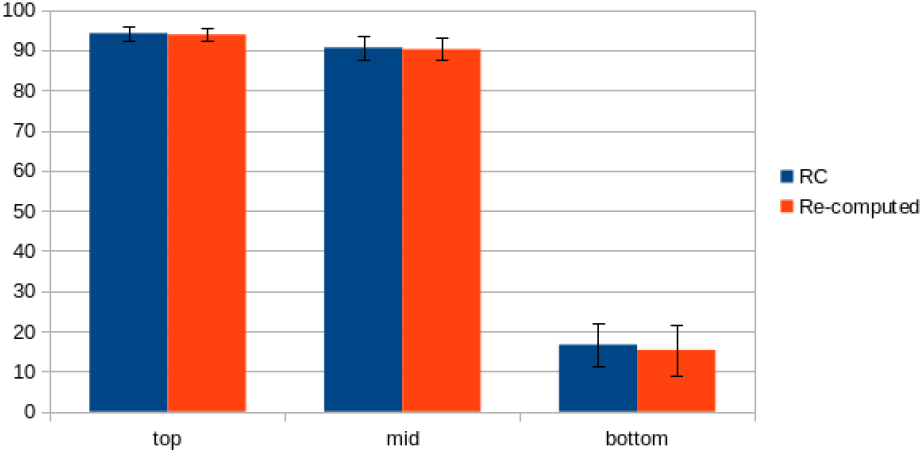
Percentage of genes from the RNA-seq related to the networks found in the resulting networks. The RNA-seq genes are separated in three categories: the top 10% most expressed, the middle 10% and the 10% least expressed. Each color represent one way of computing the tissue-specific networks, blue the original networks and red the recomputed ones. Analysis done on the 12 networks.

#### 5.2.2. Scores distributions

As presented in the previous subsection the re-computed networks have a different topology from the original study, and are portions of the original networks. But the *Regulatory Circuits* results were not only the relations but also the associated scores.

We looked at the score computed in the two versions of the networks (Figure 8): the score found in the original *Regulatory Circuits* study are in average 10 times lower that the re-computed scores.

**Figure 8:**
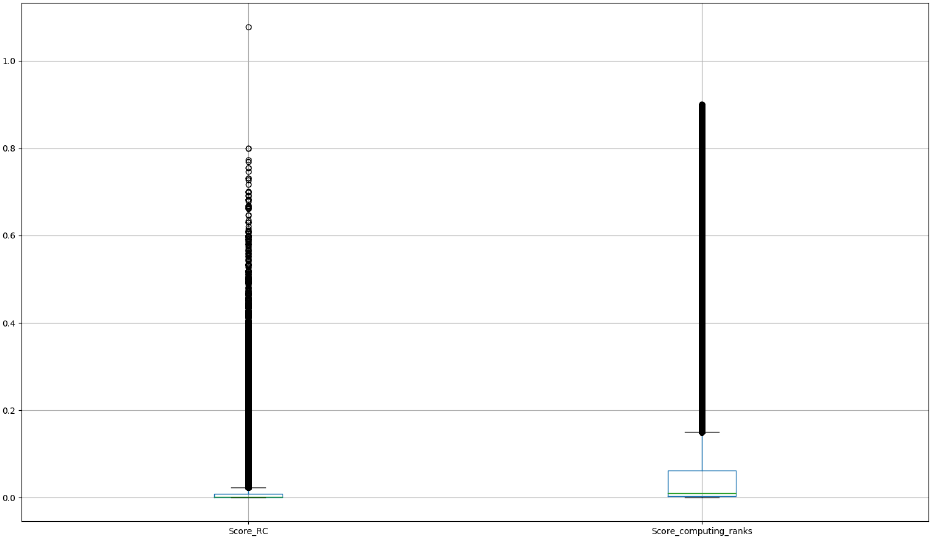
Distribution of the scores across the three methods of calculating *Regulatory Circuits* networks. Focus on B lymphoblastoid cell line. Scores in the original network: min 2.82E-7, max 1.08 and mean 0.01. Scores when re-computing the ranks: min 4.45E-6, max 0.9 and mean: 0.08

This raised the issue of the combination of the enhancer and promoter networks scores, as the original study has scores greater than 1 and our method does not. We were once again confronted to the methods section imprecision on the score computation. As the intermediary files for the networks computed by enhancers and promoters were not available, we were not able to confirm the “added” notion and how the scores are combined to produce the final networks (see 3.5). However, by inspecting the networks published by *Regulatory Circuits*, we found relations with a score greater than 1, suggesting that the final scores were obtained by adding the enhancer and promoter scores for each TF-gene relation. With our re-computed scores, performing an addition on the enhancers and promoters scores for a same TF-gene relation could have yielded a result greater than 1. However, using the addition would have resulted in even greater scores, therefore, we chose to keep the maximum of both scores, which still resulted in scores greater than *Regulatory Circuits* and was consistent with the other steps of the workflow, where the authors always used the maximum as the merging factor between redundant relations.

Since the score distribution seemed to vary between the original networks and the recomputed one, we computed the correlation between the scores. Figure 9 shows that there is a correlation between the scores in the original study and in the re-computed one (*r* > 0.5). This is also the case in the other networks observed from r=0.556 in the Brain (adult) network and r=0.785 in the Epitheloid Carcinoma network.

**Figure 9:**
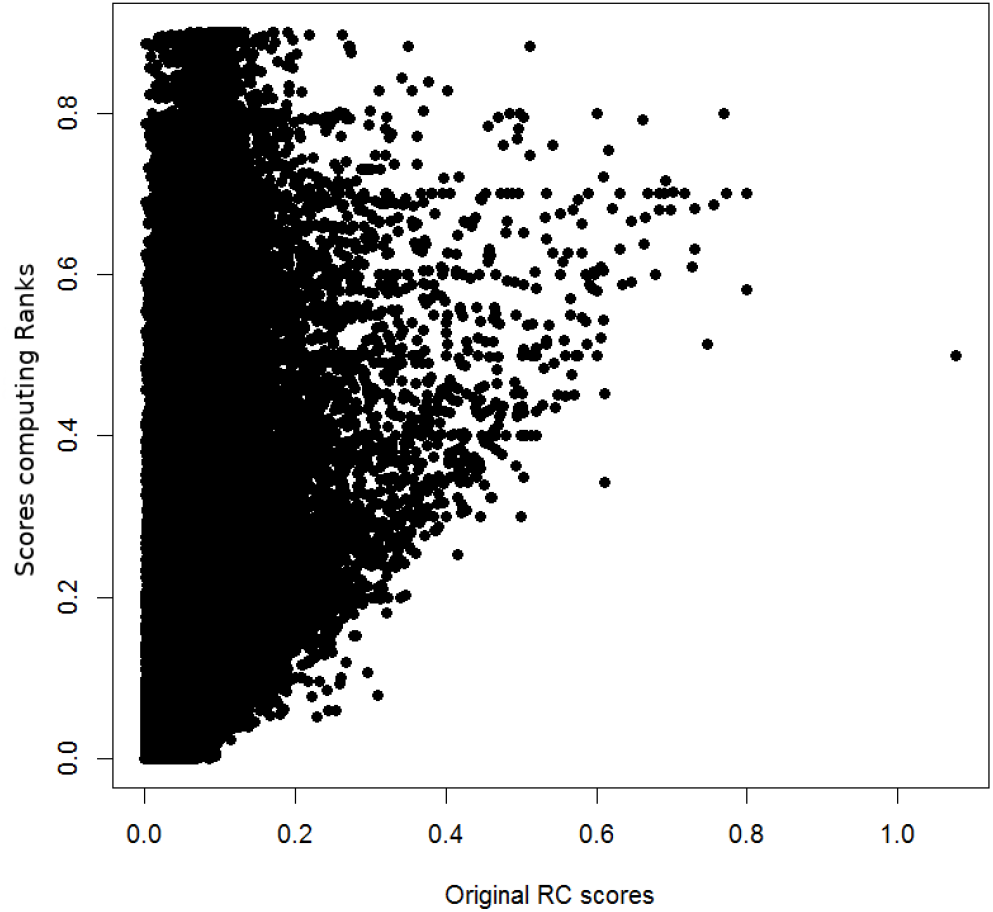
Correlation between the scores found in the original networks and the scores found in the re-computed networks for the conserved relations. Focus on B lymphoblastoid cell line. The re-computed rank networks have a r of 0.696.

For each relation, we compared its score in the re-computed network to its score in the original network of the same tissue. We found that the conserved relations have significantly (p < 2.2e-16, Wilcoxon rank sum test) higher scores than the relations excluded in the recomputed networks (see Figure 10), suggesting that conserved relations between the original *Regulatory Circuits* networks and our new computation were the most probable ones.

**Figure 10:**
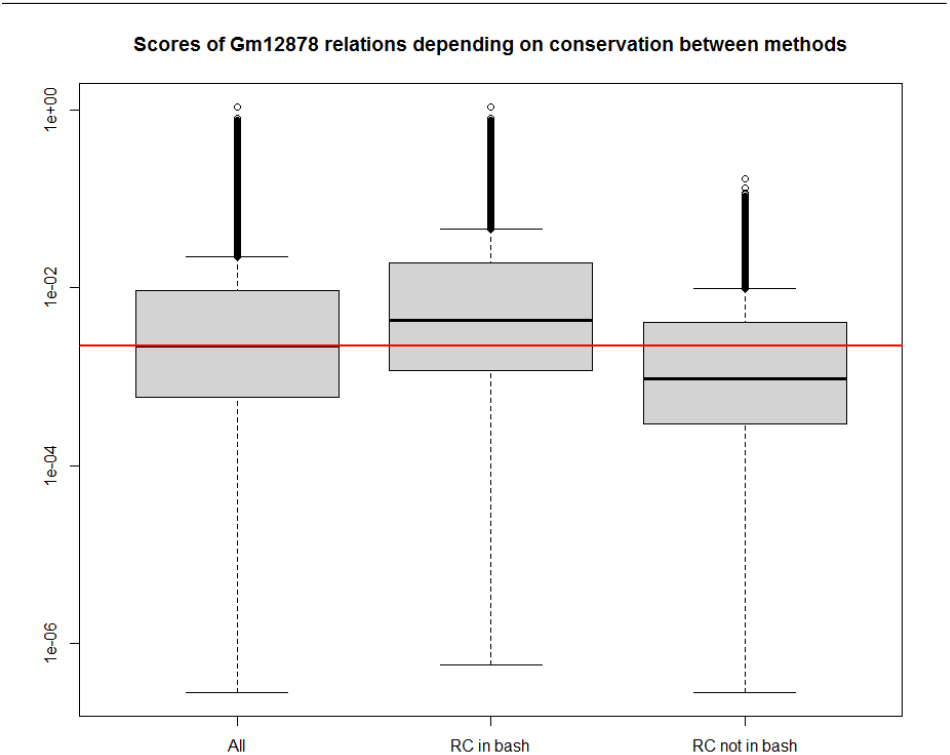
Distribution of the scores across the two methods of calculating *Regulatory Circuits* networks, depending of the conservation of the relations from the original network. Focus on B lymphoblastoid cell line. Score presented as log10 of the computed scores.

## 6. Conclusion

*Regulatory Circuits* is a major biological resources providing cell type-specific regulatory networks in Human. It has been extensively cited and reused in other studies. However, it presents several reproducibility challenges that prevent its evolution and extension, as well as reusability challenges. The technical and methodological aspects of these challenges may be also present in similar resources, which makes *Regulatory Circuits* a relevant use case for assessing reproducibility and reusability.

Our first contribution is an in-depth analysis of *Regulatory Circuits* allowing the identification of the factors limiting its reusability or reproducibility. We demonstrated how these factors prevented the direct reuse of the data set, its replication by executing the workflow on the original primary files, and its transposition by executing it on updated, extended or limited primary files. We assume that these factors are also applicable to most of the life science resources produced by an analysis workflow.

Our second contribution is an analysis of some limitations of the *Regulatory Circuits* method. They hamper reusability from a biological perspective. Firstly, using a rank function for normalization is not robust, as a noisy sample could influence the rank of the other samples. Secondly, the *Regulatory Circuits* workflow does not check the expression of TFs in the samples, meaning that some potentially absent TFs are given regulatory impact in the regulatory networks. Eventually, the activation relations are highly favored by the workflow, as the highly expressed genes and active regions have a greater weight in the score calculation, leading to poorly explained regulation of lesser expressed genes.

Our third contribution is a new implementation of the *Regulatory Circuits* workflow. This was necessary because there are eighteen java tools – across 3 folders – provided by *Regulatory Circuits* but no explanation about what each of them does, about the parameters to provide nor about the order in which they should be executed. We developed a new implementation of the complete data analysis workflow based on the available information on the methodology described in the original article. This new implementation addresses the reproducibility challenges as the dataset can now be recomputed. It revealed that 36.5% of the regulation relations of the original dataset could not be explained.

Those issues are common in life science data and impair the reproducibility and the reusability of many studies. Nonetheless, we believe that our implementation will support both reproducibility and reusability of *Regulatory Circuits*.

## ACKNOWLEDGEMENTS

We acknowledge the GenOuest bioinformatics core facility (https://www.genouest.org) for providing the computing infrastructure.

## Footnotes

1 http://regulatorycircuits.org/

2 https://github.com/marbach/magnum-app/wiki

3 http://regulatorycircuits.org/

